# Setting new boundaries of 16S rRNA gene identity for prokaryotic taxonomy

**DOI:** 10.1101/2025.02.04.636517

**Authors:** Timothy J. Hackmann

**Affiliations:** Department of Animal Science, University of California, Davis, CA, USA

**Author notes:** Correspondence: Timothy J. Hackmann.

**Keywords:** 16S rRNA gene, taxonomy, prokaryotes

## Abstract

The 16S ribosomal RNA (rRNA) gene is frequently sequenced to classify prokaryotes and identify new taxa. If sequences from two strains share less than ∼99% identity, the strains are usually classified as different species. Classification thresholds for genera and other ranks have also been proposed, but they are based on dated datasets. Here we update these thresholds by determining sequence identity of the 16S rRNA gene for nearly 18,000 type strains. This represents 86.5% of all strains validly published, and it involved making more than 161 million pairwise sequence alignments. In 90% of all cases, sequences from the same species shared a minimum of 97.2 to 100% identity. The corresponding values for genus were 90.3 to 99.0%, 79.6 to 94.1% for family, 73.1 to 90.1% for order, 72.2 to 86.3% for class, and 69.7 to 83.6% for phylum. We propose these values serve as thresholds for classifying new prokaryotic taxa. A major change from previous guidelines is recognizing that these boundaries overlap. This overlap has already been observed for relative evolutionary divergence, a metric correlated with 16S rRNA gene identity. Together with other metrics, 16S rRNA gene identity allows classification of prokaryotes from species to phylum.

## INTRODUCTION

Sequencing the 16S ribosomal RNA (rRNA) gene is a fast and versatile way to classify prokaryotes and identify new taxa. The gene is present in all prokaryotes, easily amplified by PCR [1], and already sequenced for many type strains. A sequence from a new strain can be compared to others using web servers [2, 3] or other approaches. If the sequence shares less than ∼99% identity with existing ones, the strain is usually classified as a new species [4, 5]. Similar thresholds for identity have been proposed for other ranks [6], allowing classification of strains from species to phylum.

Reliable classification depends on using reliable thresholds for sequence identity. The most widely-used thresholds were set more than 10 years ago [4–6]. With changes to taxonomy and availability of more sequences, it is unclear if these older thresholds are still reliable. Our objective was to update these thresholds using a dataset of 18,000 prokaryotic type strains. This represents all strains available with appropriate information, and our analysis revealed new boundaries for classification.

## METHODS

### Overview

Unless otherwise noted, procedures were done using R statistical software. Some functions were written in C++ and called in R using the Rcpp package. Code was run on a workstation with an Intel Core i7-14700 vPro processor and 64 Gb RAM.

### Downloading data

Taxonomy of strains was downloaded from the List of Prokaryotic names with Standing in Nomenclature (LPSN) database [7]. We downloaded the list of genera, species, and subspecies from https://lpsn.dsmz.de/downloads. The date of download was 17 October 2024. We kept all strains that had “status” containing “correct name”. To add higher ranks to this list, we downloaded *.html pages for each strain and their parent taxa. We used the polite package of R for downloading, following rules on the robots.txt file of LPSN’s website. From the downloaded pages, we used the rvest package and html selectors to extract names of ranks.

Sequences were downloaded from the LPSN database using a similar approach. From the *.html pages previously downloaded, we used the rvest package of R and html selectors to extract links to fasta files. We then used the httr package to perform downloading and BioStrings packages to read and save fasta files. We combined taxonomy and sequences into a single dataset (Data S1). This dataset had a total of *n* = 20,806 strains and *n* = 20,278 sequences.

Data from the Genome Taxonomy Database (GTDB) [8] were downloaded from https://data.ace.uq.edu.au/public/gtdb/data/releases/latest/. The date of the download was 11 November 2024. These data included a phylogenetic tree (bac120.tree) and metadata (bac120_metadata.tsv.gz). We kept all strains that had “gtdb_type_designation_ncbi_taxa_sources” equal to “LPSN” and “gtdb_representative” equal to “TRUE”. All data used were for bacteria. We did not use data for archaea because few strains (*n* = 13) were shared with and the same taxonomy as in the LPSN.

### Filtering of sequences and taxonomy

Sequences from the LPSN were filtered for length and quality. We removed any sequences shorter than 1,100 or longer than 1,750 characters. We also removed sequences with more than 15 ambiguous characters (not A, T, C, or G). A total of *n* =2,745 sequences (13%) were removed.

After filtering sequences, we removed strains with ambiguous taxonomy. These strains were those with taxonomic ranks containing “No-Family”, “No-Order”, or “No-Class”. A total of 67 strains (0.4%) were removed, leaving *n* = 17,994 strains with sequences for the analysis.

### Pairwise alignment and sequence identity

We performed pairwise alignment of sequences with the RSeqAn package of R. This package calls functions of the SeqAn C++ library [9]. To perform alignment, we used the globalAlignment() function with match score of 2, mismatch score of -1, gap open score of -10, and gap extend score of -0.5.

We calculated pairwise identity of sequences using C++ functions. The formula was identity = matching characters/total characters in alignment * 100 Following EzBioCloud [2], we did not count characters in positions with gaps. The calculation was performed in parallel by splitting data (sequence pairs) into 1,000 chunks, then processing chunks with the future [10] and furrr packages of R.

### Use of EzBioCloud

We ran *n* = 100 sequences as queries in EzBioCloud [2], a web server for strain identification. After running the query, we selected the “Valid names only” option, and then downloaded results as Excel files. We compared values of identity (similarity) from EzBioCloud to those in our analysis, matching strains according to genus and species names.

### Relative evolutionary divergence

We calculated values of relative evolutionary divergence (RED) [11] for ancestors of strains. The phylogenetic tree from GTDB [8] was used in all analyses. With this tree, we found the node of the most recent common ancestor (MRCA) for a given pair of strains. The node was found using a C++ function. We then found the value of RED for that node with the get_reds() function of the castor package [12] of R. We tested these functions using a tree with known values of RED (Fig. 1 in ref. [11]). The calculation was performed in parallel after splitting data (strain pairs) into chunks.

**Fig 1.**
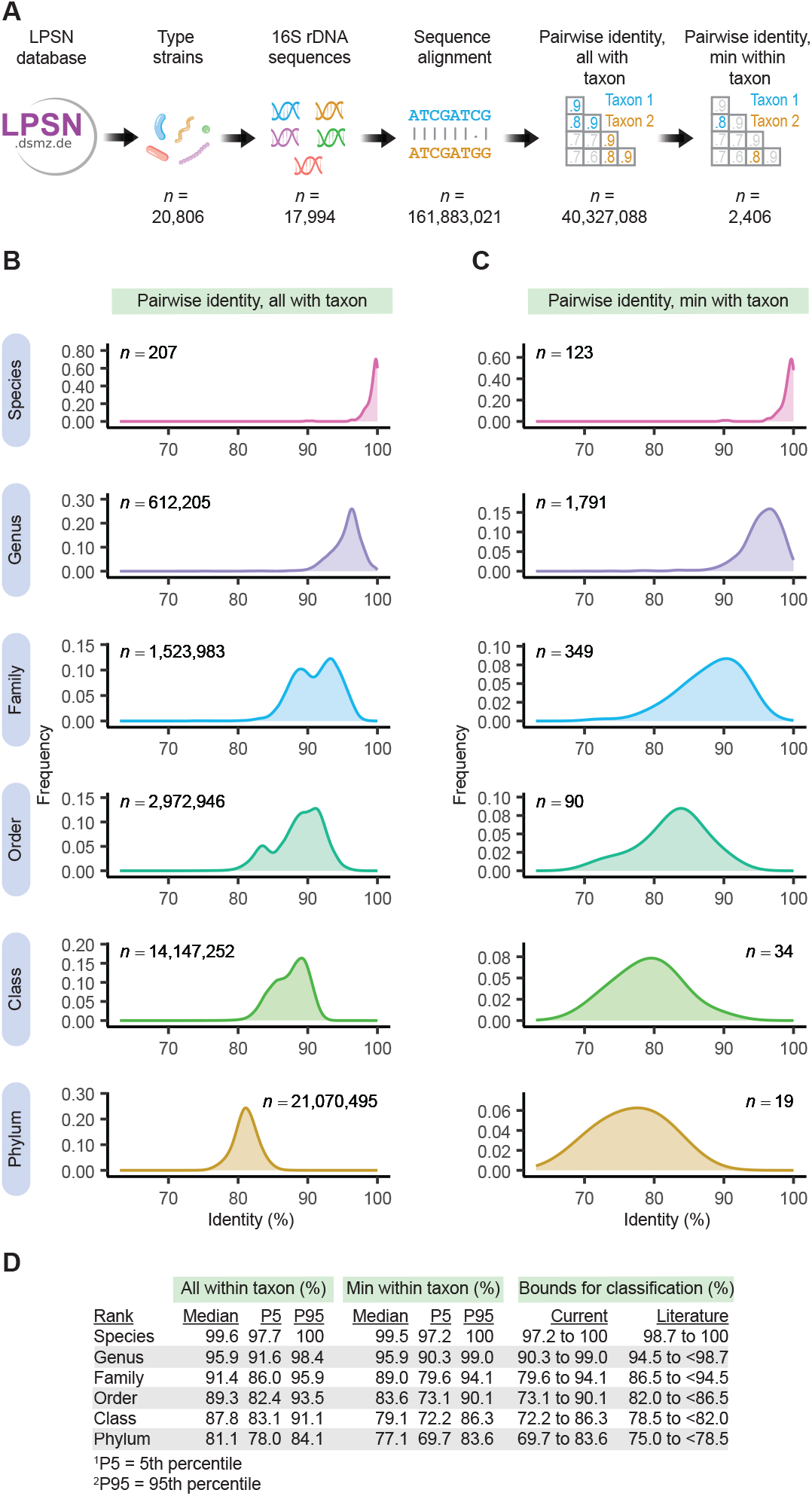
Analysis of 16S rRNA gene sequences from type strains shows values of sequence identity within taxonomic ranks. (A) Workflow for analysis. (B) All values of pairwise identity within a given taxon. (C) Minimum values of pairwise identity within a given taxon. (D) Summary and recommended boundaries for classification. Values in the literature are from ref. [4] for species and ref. [6] for all other ranks.

### Analysis of results

We calculated summary statistics with functions of the dplyr package of R. Figures were plotted using the ggplot2 package [13]. In density plots, kernel density estimation was done with the geom_density() function with the adjust parameter set to 1.5. In scatterplots, local regression was done with the geom_smooth() function and the loess method. Due to their large size, datasets were loaded using functions of the Arrow and DuckDB packages before analysis.

## RESULTS

Older literature suggests that prokaryotes are different species if their 16S rRNA sequences share less than ∼99% identity [4, 5]. To determine if this and similar thresholds [6] are still reliable, we analyzed 16S rRNA sequences from all type strains from LPSN database [7]. This database contains all species that have been validly published in the literature, along with their taxonomy and 16S rRNA gene sequences.

Our analysis involved nearly 18,000 sequences and more than 161 million pairwise sequence alignments (Fig 1A). It showed values of pairwise identity vary within taxonomic rank (Fig. 1B,C). Values fall in the order expected, being highest for species and lowest for phyla, with overlap across ranks. This is observed whether all values are considered (Fig. 1B) or only the minimum value within a given taxon (Fig. 1C). The latter type of values (minimum within taxon) have been used to set thresholds for classification in previous work [6]. We therefore summarized values in our analysis and used them to set new thresholds for classification (Fig. 1D, Data S2). The upper and lower boundaries for classification correspond to 5^th^ and 95^th^ percentiles of values in our analysis. Compared to thresholds in the literature [4, 6], our recommended boundaries for classification are lower and broader.

Using 5^th^ and 95^th^ percentiles as boundaries helps exclude outliers, which are present in our analysis. One outlier occurs with members of *Syntrophomonas wolfei* species. These members share only 90.6% identity (Data S2), falling well below the 97.2% boundary for species (Fig. 1D). This low value of identity has been recognized and led to a proposal that these members be assigned to different species [14]. Excluding these outliers makes our boundaries more reliable for classification, though the outliers themselves deserve further study.

We analyzed 16S rRNA sequences with custom R and C++ scripts, but many users perform this analysis with web servers instead. This motivated us to compare values from our analysis to those of EzBioCloud [2], a web server. We chose *n* = 100 sequences our analysis at random, ran them as queries in EzBioCloud, and recorded values of identity of hits. We found values of identity from EzBioCloud closely matched those in our own analysis (Fig. S1). This supports our approach for analyzing these sequences.

Our analyses have focused on sequence identity, but another way to classify prokaryotes is with relative evolutionary divergence (RED) [11]. We thus did another analysis to compare sequence identity to RED (Fig. 2A). We used all type strains shared by LPSN [7] and GTDB [8], the database that introduced RED as a classification metric. We considered only strains with the same taxonomy (from phylum to species) in both databases. When plotted with rank, values of identity and RED showed a similar pattern, being highest for genus and lowest for phylum (Fig. 2B,C). Values of each metric overlapped across ranks, though RED had less overlap than sequence identity at high ranks (e.g., class and phylum). When plotted against each other, values of identity and RED were correlated, with the correlation being strongest at high values of each metric (Fig. 2D). Because GTDB has only one representative strain per species, our analysis did not include strains belonging to the same species. These results show that RED and identity give similar, but not identical, results.

**Fig 2.**
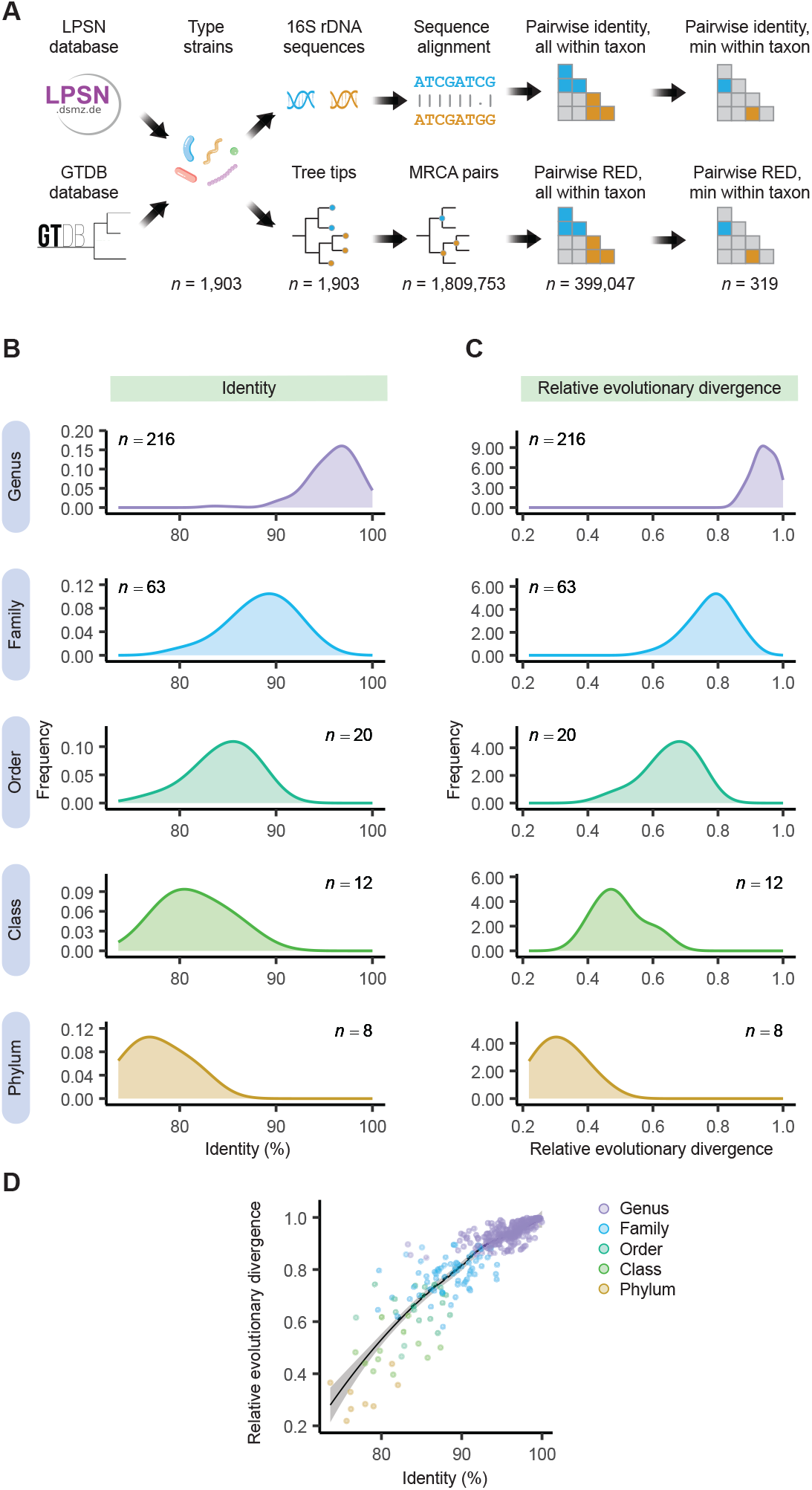
Analysis of type strains shows 16S rRNA gene identity is correlated with relative evolutionary divergence (RED), another metric used for classifying prokaryotes. (A) Workflow for analysis. (B) Minimum values of pairwise identity within a given taxon. (C) Minimum values of RED within a given taxon. (D) Values of identity vs. RED.

In sum, our study shows values of 16S rRNA gene identity expected within and across ranks. It shows values overlap across ranks, and it proposes overlapping boundaries for classification in turn. Values of identity were correlated with RED, showing they could be complementary in classifying prokaryotes.

## DISCUSSION

After isolating a new prokaryotic strain, it is routine to sequence its 16S rRNA gene and determine its identity to sequences of other strains. If the identity is low enough, the strain is classified as belonging to a new species or another taxonomic rank. While this approach is common practice, it is only reliable if there are reliable thresholds used for classification. Thresholds in the literature were set years ago [4–6]. An additional consideration is some thresholds were set indirectly—by comparing sequence identity to another metric of genetic relatedness [4, 5]. Another threshold was set using an unclear dataset (no strain names reported) [15]. Reexamination of these thresholds is overdue.

Our analysis uses sequences from 18,000 type strains, more than doubling the number used in the past [5, 6]. It also uses taxonomy that is up-to-date and with validly published names only [7]. Our results show the 16S rRNA gene is useful for classification, but simple thresholds (e.g., 94.5% for genus) need to be reconsidered. Because identities overlap across ranks, we suggest that classification be done using boundaries that overlap, also. Under this approach, a strain sharing 94% identity with another strain could be classified as belonging a new genus, but it could also be classified as belonging to a new family. Allowing overlap is a change but is already practiced with RED [11], another metric for classifying prokaryotes.

Prokaryotes are increasingly classified using whole-genome sequences [16], not 16S rRNA gene sequences alone. Relative evolutionary divergence (RED) is one such metric that depends on whole-genome sequences [11], and it has been used to classify prokaryotes from genus to phylum [17]. Our results show that both RED and sequence identity can be used for classification. Sequence identity is more useful at low ranks—it can classify strains up to the level of species— while RED can be more discriminatory at high ranks. Thus, their use is complementary. Other metrics depending on whole-genome sequences include average nucleotide identity (ANI) [18, 19], average amino acid identity (AAI) [20], percentage of conserved proteins (POCP) [21], and digital DNA–DNA hybridization (dDDH) [22]. These metrics are useful at low ranks, but those tested do not discriminate well at ranks above species [5] or genus [16, 20].

Our analysis relies on type strains and a regulated taxonomy, presenting both strengths and weaknesses. Type strains are well characterized, but they include only cultured organisms. Most prokaryotes are uncultured [23], and their genome sequences are often missing 16S rRNA genes. This points to a broader need for more cultured organisms and more complete genome sequences. The taxonomy we use contains validly published names [7], but that taxonomy can still have inconsistencies. One inconsistency is with members of *Syntrophomonas wolfei*, which differ genetically and phenotypically [14] yet are still considered a single species. Our boundaries for classification were set to exclude the worst outliers, though inconsistences are still invariably present.

In sum, our study provides practical boundaries for using the 16S rRNA gene to classify prokaryotes from species to phylum. By allowing boundaries to overlap between ranks, it provides a framework for classification in line with our current system of taxonomy. Compared to other approaches for classification, sequencing the 16S rRNA gene is a simple, fast, and versatile option. It has been routinely used for over three decades [24], and we hope our recommendations will support its use for decades to come.

## DATA AVAILABILITY

Code for downloading data from the LPSN, performing pairwise alignments, calculating sequence identity, and determining relative evolutionary divergence is available at https://github.com/thackmann/SequenceIdentity.

## AUTHOR CONTRIBUTIONS

T.J.H.: Writing - Review & Editing

