## Supplementary Material for "Setting new boundaries of 16S rRNA gene identity for prokaryotic taxonomy"

Timothy J. Hackmann*

**This PDF file includes:**

Figure S1
Legends for Data S1 to S2

**Other Supplemental Material for this manuscript includes the following:**

Data S1 and S2

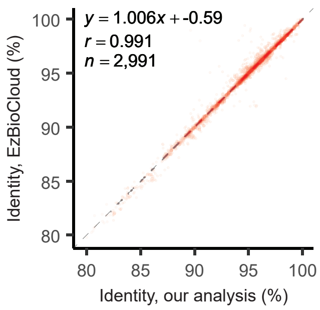


**Fig. S1.** Pairwise identity calculated in our analysis closely matches that of EzBioCloud.

**Data S1. (separate file)**
Taxonomy and sequences of strains used in our analysis.

**Data S2. (separate file)**
Values pairwise identity summarized by taxon.
